# Imprint of Assortative Mating on the Human Genome

**DOI:** 10.1101/300020

**Authors:** Loic Yengo, Matthew R. Robinson, Matthew C. Keller, Kathryn E. Kemper, Yuanhao Yang, Maciej Trzaskowski, Jacob Gratten, Patrick Turley, David Cesarini, Daniel J. Benjamin, Naomi R. Wray, Michael E. Goddard, Jian Yang, Peter M. Visscher

## Abstract

Non-random mate-choice with respect to complex traits is widely observed in humans, but whether this reflects true phenotypic assortment, environment (social homogamy) or convergence after choosing a partner is not known. Understanding the causes of mate choice is important, because assortative mating (AM) if based upon heritable traits, has genetic and evolutionary consequences. AM is predicted under Fisher’s classical theory^1^ to induce a signature in the genome at trait-associated loci that can be detected and quantified. Here, we develop and apply a method to quantify AM on a specific trait by estimating the correlation (*θ*) between genetic predictors of the trait from SNPs on odd versus even chromosomes. We show by theory and simulation that the effect of AM can be distinguished from population stratification. We applied this approach to 32 complex traits and diseases using SNP data from ∼400,000 unrelated individuals of European ancestry. We found significant evidence of AM for height (*θ*=3.2%) and educational attainment (*θ*=2.7%), both consistent with theoretical predictions. Overall, our results imply that AM involves multiple traits, affects the genomic architecture of loci that are associated with these traits and that the consequence of mate choice can be detected from a random sample of genomes.

---

Non-random mating in natural populations has short and long-term evolutionary consequences. In many species, including humans, mate choice is often associated with phenotypic similarities between mates^2,3^. Such phenotypic similarities have multiple sources, for example social homogamy, the preference for a mate from the same environment, or because of primary assortment on certain traits observable at the time of mate choice. Contrary to social homogamy, primary phenotypic assortment, here referred to as assortative mating (AM), has genetic and evolutionary consequences and therefore is the focus of our study. In humans, AM involves multiple complex traits^4–8^ and can sometimes lead to similar susceptibility to diseases9–12. The genetic effects of AM were first studied in the seminal articles of Fisher (1918)^1^ and Wright (1921)^13^. Those two founding contributions, further complemented by Crow & Kimura (1970)^14^ and others^15–17^ have set the basis of the theory of AM on complex traits. AM theory predicts three main genetic consequences of a positive correlation between the phenotypes of mates in a population: (i) an increase of the genetic variance in the population, (ii) an increase in the correlation between relatives and (iii) an increase of homozygosity (deviation from Hardy-Weinberg Equilibrium; HWE), in particular at causal loci. These seemingly distinct consequences are nonetheless explained by the same cause: directional correlation between trait-increasing alleles (TIA), also referred to as gametic phase disequilibrium (GPD), induced both within and between loci (**Fig. 1**). AM-induced GPD implies correlations between physically distant loci (between chromosomes for example) and is thus distinct from linkage disequilibrium. Therefore, AM leads to a genomic signature of trait-associated loci that can be quantified by estimating GPD.

**Fig. 1.**
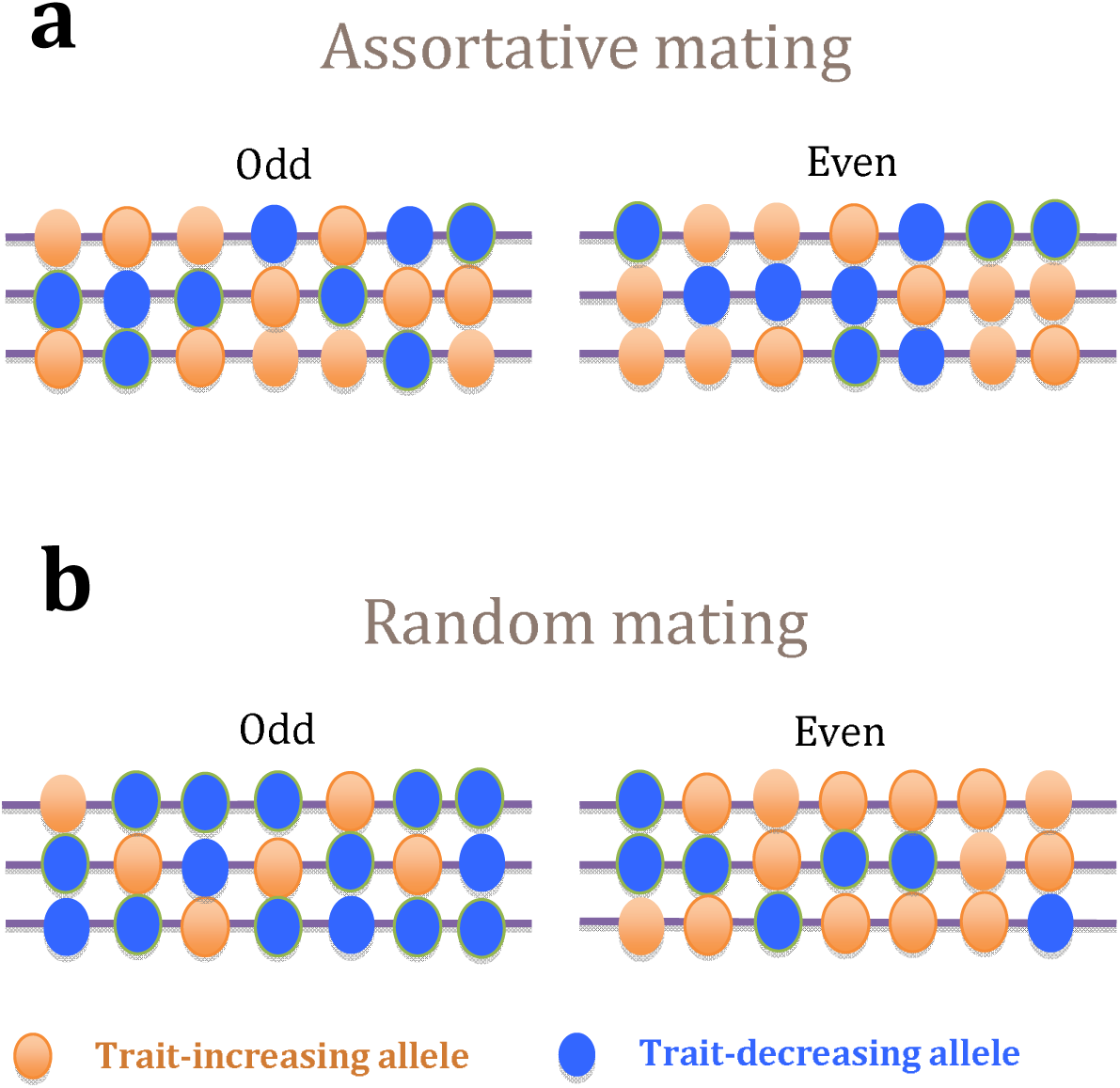
Schematic illustration of the effect of assortative mating (AM) on the correlation between trait-associated alleles. Each line represents a chromosome of an individual in the population and each coloured bead represents an allele (orange: trait-increasing alleles (TIA); blue: trait-decreasing alleles) at a particular locus on that chromosome. Panel **a** represents a population undergoing AM and panel **b** represents a population undergoing RM. Under RM the distribution of alleles between odd and even chromosomes are uncorrelated (no-consistent pattern between chromosomes). Under AM, the distributions of alleles are correlated between chromosomes, such that the number of TIAs on odd chromosomes predicts the number of TIAs on even chromosomes.

Previous studies^18–20^ have been successful at detecting GPD by direct quantification of increased homozygosity at ancestry-associated loci. Beyond ancestry-related traits, such endeavour is particularly challenging for polygenic traits as theory^14^ predicts an increase of homozygosity due to AM inversely proportional to the number *M* of causal variants^14,21^. For a highly polygenic trait like height with an estimated *M*∼100,000 for common variants^22^, the expected increase in homozygosity would be of the order of ∼1/2*M* =5×10^−6^, i.e. negligible (**Supplementary Note 1**). Extremely large studies would therefore be required to quantify systematic deviation from HWE at height-associated single-nucleotide polymorphism (SNP) as shown in a recent study^18^ that failed to detect such an effect. Another study^23^ in ∼6,800 participants of European ancestry, reported evidence of deviation from HWE at height associated loci. This study however did not account for within-sample population stratification and therefore their reported estimates are likely inflated for this reason. Overall, study designs using deviation from HWE for quantifying GPD can be successful for detecting ancestry-based AM (ancestral endogamy) because the number of loci distinguishing ancestries is relatively small^24^, and ancestral endogamy is strong^18^, but are less powerful to detect trait-specific AM. In contrast to HWE-based estimation strategies, quantifying GPD on the basis of pairwise correlations between TIAs is much more tractable as the number of pairs of loci involved, of the order of ∼*M*^2^, compensates for the magnitude of the expected covariance for each pair, ∼1/2*M*. This compensation explains in essence why AM increases the genetic variance in a population^14,21^.

GPD due to AM causes individuals that carry TIAs at one locus to be more likely to carry TIAs at other loci than expected under gametic phase equilibrium. Consequently, individuals with many TIAs on even chromosomes are likely to have more than average TIAs on odd chromosomes. We quantify this effect by calculating genetic predictors for a trait from the SNPs on odd chromosomes and from the SNPs on even chromosomes and then calculating the correlation (*θ*) between these two predictors. To calculate these predictors we use estimates of the effect of each SNP on a trait from publicly available summary statistics from genome-wide association studies (GWAS) of large sample size. We apply these estimated SNP effects to individuals in a separate sample who have SNP genotypes available. We can calculate the trait predictor based on odd and even chromosomes separately and estimate the correlation between them (i.e. *θ*). Our method measures the effect on the genome due to AM in previous generations and thus does not require observed phenotypes or the use of mate pairs. Under the null hypothesis of random mating (RM), the correlation between alleles on different chromosomes is expected to be 0 as a consequence of the independent segregation of chromosomes. However, population stratification can induce spurious correlations between alleles, even at distant loci. Intuitively, if *θ* is estimated from a mixture of two sub-populations with distinct allele frequencies, then having TIAs more frequent in one of the sub-populations (even by chance) would result in an apparent correlation between TIAs even when such correlation is absent in each sub-population (**Supplementary Note 2**). We show through simulations how the effect of population stratification can be corrected with our method. We then applied our method to estimate AM-induced GPD for 32 traits and diseases in samples of unrelated genomes from three independent cohorts: ∼350,000 participants of the UK Biobank (UKB), ∼54,000 participants of the Genetic Epidemiology Research on Adult Health and Aging (GERA) cohort and ∼8,500 participants of the US Health and Retirement Study (HRS). We find evidence of AM for a number of complex traits, including height and educational attainment.

## Results

### Theory underlying the estimation of GPD from SNP data

We derived (**Supplementary Note 1**) the expected value of the correlation across individuals between the trait predictors from SNPs on odd (*S_o_*) and even (*S_e_*) chromosomes as a function of the phenotypic correlation between mates (*r*), the equilibrium heritability of the trait 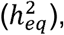
the fraction of that heritability captured by SNPs (*f_eq_*), the sample size (*N*) of the reference GWAS in which effect sizes are estimated using classical linear regression, one SNP at a time; and the number of causal loci (*M*) contributing to the trait (which differs from the number of statistically associated loci). The main result is that for a large number of trait loci,

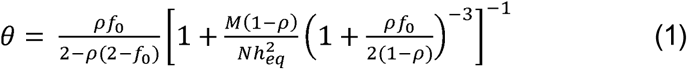

with
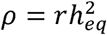
being the correlation between additive genetic values of mates expected under AM^17^ and *f_o_* ≈ *f_eq_*/( 1– *ρ*), the fraction of heritability captured by SNPs in the base population (**Supplementary Note 1**). These parameters do not need to be known or estimated, but can be used to provide *a priori* expectation of *θ* or *a posteriori* rationalisation. Hence, quantification of GPD can be directly obtained from estimates of *θ* using empirical data. For the sake of simplicity, we hereafter refer to estimates of *θ* as estimates of GPD. Equation (1) implies that the expected correlation *θ* between *S_o_* and *S_e_* increases with *N*, i.e. with better estimation of SNP effects from the reference GWAS, and with *f_eq_*, i.e. with better tagging of causal variants underlying the full narrow sense heritability.

We derived (**Supplementary Note 2**) that estimates of *θ* from the regression of *S_o_* on *S_e_* can be inflated by population stratification, especially when TIAs are highly differentiated between sub-populations. We performed a number of simulations (**Supplementary Notes 2 and 3**) to validate the impact of population stratification on our estimator of GPD and show how to adjust for it using principal components derived separately for odd and even chromosomes (**Fig. S1** and **Fig. S2**). We used this strategy to quantify GPD in real data, and therefore adjusted all GPD estimates for 20 genotypic principal components to correct within sample population stratification (**Materials and Methods**). Also, given that most GPD estimates are small, all GPD estimates (correlations) reported below are expressed as percentages (e.g. 3% instead of 0.03).

### Quantifying AM-induced GPD in complex traits

We first analysed height and educational attainment (EA), two reference traits with long-standing evidence of a positive correlation between mates. For height, we used estimated effect sizes, from summary statistics of the latest GWAS meta-analysis of the GIANT consortium (Wood et al. 2014)^25^, of 9,447 near independent HapMap3 (HM3) SNPs selected using the LD clumping algorithm implemented in the software PLINK^26^ (linkage disequilibrium (LD) r^2^<0.1 for SNPs < 1 Mb apart and association *p*-value < 0.005). We thus selected these SNPs to be enriched for true association with height. We estimated in UKB participants the correlation between height increasing alleles on odd versus even chromosomes to be *θ*_height_=3.0% (s.e. 0.2%; **Fig. 2**) and replicated this finding in GERA (*θ*_height_=4.1%, s.e. 0.4%; **Fig. 2**) and HRS (*θ*_height_=4.4%, s.e. 1.1%; **Fig. 2**). A meta-analysis of these three estimates yields a combined GPD among height-increasing alleles of 3.2% (s.e. 0.2%, *p*=6.5×10^−89^). To dismiss possible biases due to cryptic sample overlap or residual population stratification in summary statistics from the Wood *et al*. study, we re-estimated *θ*_height_ using summary statistics of a family-based GWAS that provide stringent control for population stratification^27^. We therefore meta-analysed summary statistics from the Robinson et al. (2015) study^27^ in 17,500 quasi-independent sib pairs with that from a similar analysis performed in 21,783 quasi-independent sib-pairs identified in the UKB (**Materials and Methods**). Using effect sizes of the 9,447 selected SNPs, re-estimated in the combined family-based GWAS, we found consistent GPD estimates between UKB not including sib-pairs (*θ*_height_=2.1%, s.e. 0.2%; *p*=8.4×10^−36^), GERA (*θ*_height_=2.1%, s.e. 0.4%; *p*=1.4×10^−6^) and HRS (*θ*_height_=2.5%, s.e. 1.1%; *p*=0.02). The meta-analysis of these three estimates yields *θ*_height_=2.1% (s.e. 0.2%; *p*=4.7×10^−42^). Note that lower estimates (2.1% vs 3.4%) are expected here because of the smaller sample size (*N*=39,283) of this family-based GWAS, as predicted by equation (1).

**Fig. 2.**
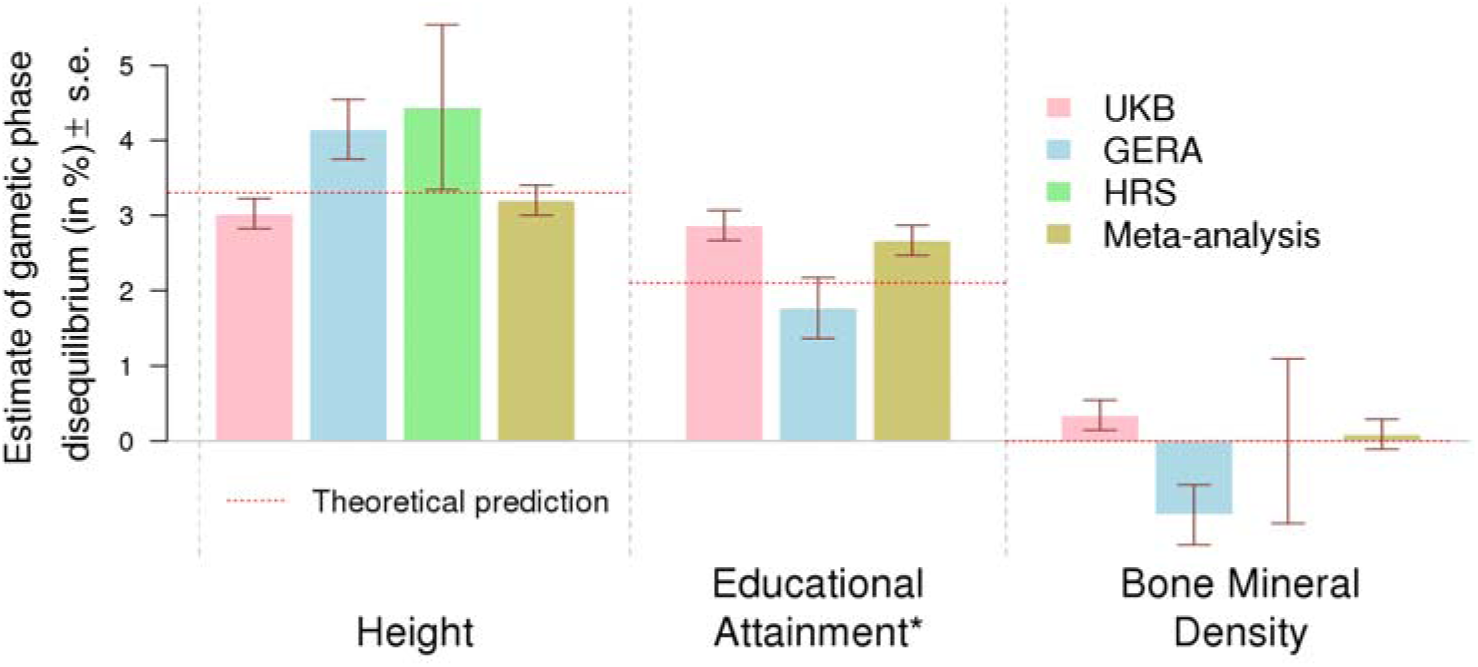
Estimate of assortative mating (AM) induced gametic phase disequilibrium (GPD) among trait increasing alleles in three independent cohorts: UKB (N=348,502), GERA (N=53,991) and HRS (N=8,552). GPD is estimated as the correlation between trait-specific genetic predictors from SNPs on odd chromosomes versus even chromosomes. Bone mineral density was selected as a trait on which AM does not occur (negative control). Estimates are adjusted for 20 genotypic principal components from SNPs on either odd or even chromosomes to correct the effect of population stratification. ^∗^The HRS cohort was not included in the meta-analysis of GPD estimates among educational attainment increasing alleles, as HRS was included in the Okbay *et al*. study. Theoretical predictions are obtained from equation (1).

For EA, we used estimated effect sizes, from the summary statistics of a large GWAS meta-analysis of the number of years of education (Okbay *et al*. 2016)^28^, of 4,618 near independent HM3 SNPs selected using the same LD clumping procedure described above. Using genotypes of 238,193 UKB participants not included in the Okbay *et al*. study (**Materials and Methods**), we found that *θ*_EA_=2.9% (s.e. 0.2%; **Fig. 2**) and replicated this finding in GERA (*θ*_EA_=1.8%, s.e. 0.4%; **Fig. 2**). We also attempted replication in HRS but the estimate we found (*θ*_EA_=8.9%, s.e. 1.1%; **Fig. 2**) was likely inflated given that HRS was part of the Okbay *et al*. meta-analysis (**Supplementary note 4**). We therefore only meta-analysed GPD estimates from UKB and GERA and found the average correlation between EA increasing alleles on odd versus even chromosomes to be *θ*_EA_=2.7% (s.e. 0.3%, *p*=6×10^−46^; **Fig. 2**). We also re-estimated the effect sizes of the 4,618 selected SNPs on EA, using the same within-family procedure described above. We found GPD estimates of ∼0.4% (s.e. 0.4%) in GERA and ∼0.3% (s.e. 0.1%) in UKB participants unrelated with any of the 21,783 sib-pairs used to estimate effect sizes. The meta-analysis of the latter two estimates is *θ*_EA_=0.31% (s.e. 0.16%; *p*=0.05). As shown below, this lower estimate is expected as the consequence of the smaller sample size used to estimate SNPs effects.

We compared GPD estimates with theoretical predictions of *θ* from equation (1). Equation (1) predicts *θ* from the sample size of the reference GWAS (N=253,288 for height and 293,723 for EA), the correlation between mates, the equilibrium heritability (80% and 40% for height and EA respectively^29^), the number of causal variants SNPs (*M*∼100,000 assumed for both traits) and the heritability captured by SNPs used to estimate *θ*. Using ∼1,000 unrelated trios (two parents and one offspring) from UKB^30^ we estimated the correlation between mates for height and EA to be 0.24 (s.e. 0.03) and 0.35 (s.e. 0.03), respectively. We estimated the SNP heritability captured by each set of SNPs used to estimate *θ* in HRS using the software GCTA^31^, resulting in estimates of
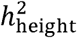
= 27.3% (s.e. 1.7%) and
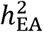
= 15.1% (s.e. 1.3%). With these five input parameters, equation (1) predicts *θ* to be ∼3.2% for height and ∼1.9% for EA. Using estimated effective sample sizes of within-family GWAS (*N_eff_* = 39,283 for height and 15,559 for EA; **Materials and Methods**), equation (1) predicts *θ* to be ∼1.3% for height and ∼0.2% for EA. Our estimates of GPD among trait-associated (*θ*_height_=3.2%, s.e. 0.2; *θ_EA_*=2.7%, s.e. 0.3%) are therefore consistent with these predictions. Everything held constant, equation (1) also predicts that with a much larger sample size of the discovery GWAS, for instance >1,000,000 participants, *θ*_height_ would be ∼4% and *θ*_EA_∼3%.

We extended our primary analysis of height and EA to detect GPD in 30 additional complex traits and diseases (**Table S1**) using the same strategy. Among these traits, we analysed bone mineral density (BMD)^32^ as a null trait given that non-significant mate correlation was previously reported^33^. As expected, we did not find significant GPD associated with BMD (meta-analysis of UKB, GERA and HRS: *θ*_BMD_=0.09%, s.e. 0.2%; *p*=0.64). After Bonferroni correction applied to the meta-analysis of UKB, GERA and HRS (*p*<0.05/32≈1.56×10^−3^), we did not detect significant GPD for any of these other traits. We believe that this observation is likely explained by lack of statistical power, in particular resulting from the smaller sizes of GWAS used for these traits (on average ∼73,000 participants compared to ∼273,000 on average for height and EA) or from smaller variance explained by SNPs selected to calculate genetic predictors of these traits. As an example, although the GWAS of body mass index (BMI) used in this study is similar in size with that of height (**Table S3**), our estimation in HRS participants of the variance explained by the 2,362 SNPs BMI-associated SNPs selected (**Table S1**) is only ∼6.2% (s.e. 0.9%) versus ∼27.3% (s.e. 1.7%) for height. A much larger GWAS would therefore be required to detect any GPD among BMI-associated alleles using our method.

### Confirmation using mate pairs

Another experimental design to quantify the genetic effect of AM on a particular trait consists of estimating the correlation of genetic predictors of this trait between mates^33–35^. We quantified the mate correlation (*r_m_*) of genetic predictors of all 32 traits (**Table S2**) using 18,984 unrelated couples identified in the UKB (**Materials and Methods**). We found significant correlations between mates for genetic predictors of height (*r_m_*=5.9%, s.e. 0.8%, *p*=9.2×10^−14^) and EA (*r_m_*=6.1%, s.e. 0.9%, *p*=7.3×10^−11^). Across all traits, we estimated the regression slope of *r_m_* estimates onto *θ* estimates to be 1.8 (s.e. 0.2) **(Fig. 3**). Both these results are consistent with our derivation that the expected mate correlation of genetic predictors is approximately twice the expected value of *θ* (**Supplementary note 4**).

**Fig. 3.**
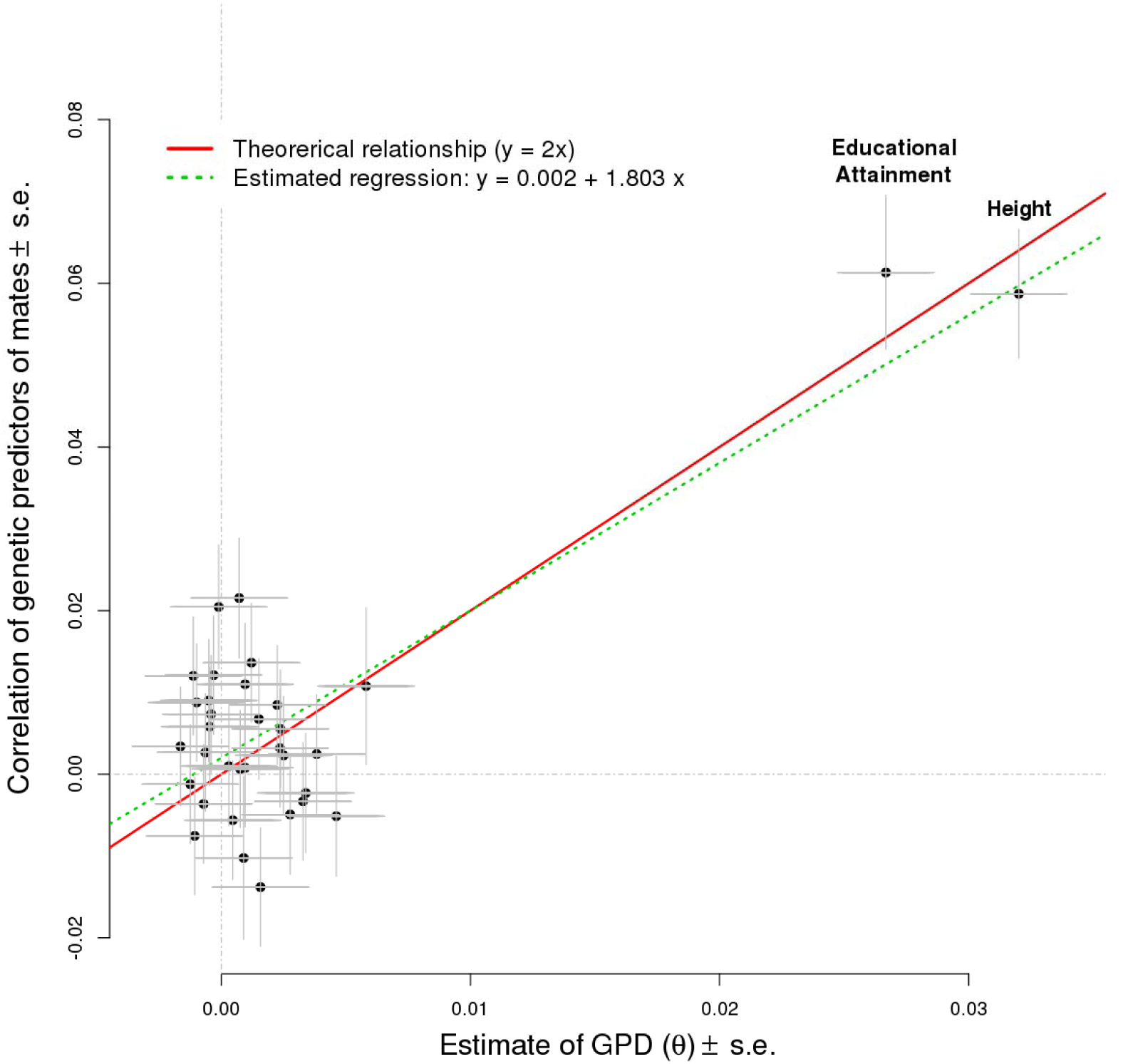
Correlation of genetic predictors in 18,984 mates pairs (y-axis; values from **Table S2**) as a function of within-individual estimates of gametic phase disequilibrium (GPD: x-axis) for 32 complex traits and diseases (meta-analysis from **Table S1** in N=411,045 participants). Theory derived in **Supplementary Note 4** predicts a regression slope equal to 2. Estimated linear regression intercept and slope are 0.002 (standard error, s.e. 0.002) and 1.8 (s.e. 0.23) respectively.

## Discussion

We have shown in this study that the genomic signature of AM can be detected and quantified using SNP data in a random sample of genomes from the population, even in the absence of data on mate pairs. This is an important aspect of our method since large datasets on mate pairs are rare and may be absent in natural populations. We confirm the genetic basis for AM for height and EA, consistent with the assumption of primary assortment on these traits. We showed that our estimates of GPD from real data are consistent with theoretical predictions made under simplifying assumptions such as equal SNP effect sizes, population being at equilibrium and that the number of common causal variants of the order of ∼100,000 (**Supplementary Note 1**). We did not however detect significant GPD for the other traits and diseases analysed in this study. Beyond true negatives such as bone mineral density, we believe that the relatively smaller size of GWAS used in our inference has reduced the power to detect the genetic signature of AM in more traits and diseases. We cannot therefore draw firm conclusion from our observations on the importance of primary assortment (as opposed to environmental correlation) to the resemblance between mates for some of these traits such as smoking habits^36^, alcohol consumption^36^ or susceptibility to psychiatric disorders^12^. Overall, our methodology is straightforward and can be applied to a wider variety of traits and in other species, provided that summaries of trait-variant associations are available. AM is multi-dimensional in essence as mate choice depends on multiple physical and behavioural traits which may or may not share a common genetic basis^5,37^. Studying networks of traits and genes driving AM is one of the challenges to meet for improving our understanding of the genetic consequences of AM in a population. As a step in this direction, our method can be for example applied to quantify consequences of AM on gene expression or at any other molecular level, through the use of SNP predictors of these endogenous traits.

Our study predicts that for traits affected by AM the estimates of SNP effects from within-family experimental designs tend to be smaller than those from a population sample, and that a genetic predictor generated from a population sample will explain less variation than expected when applied to a population not undergoing assortative mating.

Our study has a number of limitations. The first one is that certain aspects of our approach are very conservative. We have attempted to quantify GPD induced by AM while applying stringent correction for population stratification. Although such a strategy is expected to yield unbiased estimates it still faces the difficulty of distinguishing AM on population stratification related features from AM on trait specific features. Height is a typical example. AM on height occurs but, in addition, people tend to marry within geographical sub-populations (countries for example) which differ in mean height^27^. Correcting for population stratification using principal components would consequently remove part of the signal that we want to detect. We have nevertheless been able to detect GPD among height increasing alleles as a consequence of the large sample size of the discovery GWAS, the strength of assortment of mates, and the high heritability of this trait.

The second limitation relates to our strategy for SNP selection. We have included in our analyses SNPs using a low and arbitrary threshold (*p*<0.005) on the significance of association with the trait. Although this strategy is not expected to bias the covariance between *S_e_* and *S_o_*, it may increase both their variances and thus potentially induces downward biases in GPD estimates. We chose nonetheless this strategy to maximize the fraction of heritability captured by SNPs, which influences the expected correlation between *S_e_* and *S_o_* as derived in equation (1). As an example, if GPD is inferred using genome-wide significant SNPs from Okbay *et al*. (2016), which explain ∼3% of the variance of EA, the expected correlation between *S_e_* and *S_o_* would only be ∼0.45% under assumptions made above. Such small correlation is nearly undetectable in cohorts with less than 300,000 participants (**Materials and Methods**). Another SNP selection strategy could have been used to reach a better trade-off between bias and power but this would generally require observed phenotypes to optimize genetic predictors^33,34^ (find the best *p*-value threshold or shrinkage parameter).

In conclusion, our study provides empirical quantification of GPD induced among trait-associated alleles, a phenomenon predicted by theory exactly a century ago by Fisher (1918)^1^. The human genome has a pattern of trait-associated loci that is shaped by human behaviour (mate choice), as well as natural selection ^33,38–40^. The imprint of assortative mating on the contemporary human genome reflects mate choice in the 1930-1970s and in generations prior to that. Although there is much more mobility within and between human populations in the 21^st^ century, the underlying factors that determine mate choice remain stable^11,35^, and we may expect to continue to see its effect in the genome.

## Materials and Methods

### Estimation of GPD from SNP data

Our inference of GPD in a given sample of genomes is based on the correlation *θ* between polygenic scores *S_e_* and *S_o_* calculated from SNPs on even and odd chromosomes respectively. For each individual from the study population, these scores are obtained as linear combinations of SNPs allele counts weighted by their estimated effect sizes from publicly available GWAS of complex traits and diseases (**Supplementary Note 1**). We used publicly available summary statistics (regression coefficients for each tested SNP and *p*-values) from large GWAS on 32 traits (**Table S3**; URLs to download these summary statistics are given in **Supplementary Note 1**). These include GWAS on cognitive traits (educational attainment, intelligence quotient), anthropometric traits (height, body mass index, waist-to-hip ratio), psychiatric disorders (attention deficit hyperactivity disorder, autism spectrum disorder, bipolar disorder, anxiety, major depressive disorder, post-traumatic stress disorder and schizophrenia), other common diseases (coronary artery disease, type 2 diabetes, Crohn’s disease and rheumatoid arthritis), blood pressure, reproductive traits, personality traits, alcohol and smoking, and bone mineral density as a null trait. It is important that the sample of people whose genotypes were used was independent of the sample of people used to estimate SNP effects on each trait. Otherwise large biases can be expected as shown in **Supplementary Note 4**. We applied LD score regression (LDSC) for quality control and only kept summary statistics with a corresponding ratio statistic (ratio = (LDSC intercept – 1) /(mean chi-square statistic over all SNPs - 1)) non-significant from 0 (i.e. estimate / standard error < 2) or < 0.2 (**Table S3**). Significance of the GPD estimates was assessed using *p*-values from Wald-tests, with the null hypothesis “Ho: *θ*= 0” versus the alternative “H_1_: *θ*≠0”.

### Correction of population structure

We used genotypic principal components to correct for population stratification. We calculated 20 principal components from 70,531 near independent SNPs (35,301 on odd chromosomes and 35,230 on even chromosomes) selected using the LD pruning algorithm implemented in PLINK (r^2^<0.1 for SNPs < 1Mb apart). We denote these principal components as PCO for SNPs on odd chromosomes and PCE for SNPs on even chromosomes. When *θ* is estimated from the regression of *S_e_* onto *S_o_*, the effect of population stratification is corrected by adjusting the regression for PCOs (and vice versa). This can be summarised using the following regression equations: *S_e_* = *θS_0_* + PCO_1_ +…+ PCO_20_ or *S_0_* = *θS_e_* + PCE_1_ +…+ PCE_20_. Since *S_e_* and *S_o_* may not have exactly the same variance as a result of SNPs sampling, we chose to estimate *θ* from the regression onto the genetic predictor with the larger variance. We observed nonetheless that estimates obtained from the regression of *S_e_* onto *S_o_* are very similar to those obtained from regression of *S_o_* and *S_e_* (**Fig. S3**). In the simulation studies (**Supplementary Note 2**) we also considered the case where principal components are calculated from all SNPs available (odd and even chromosomes). In our simulations, principal components were calculated using R version 3.1.2, while in real data principal components were calculated using the fast PCA approach^41^ implemented in PLINK version 2.0.

### Statistical power

Theory underlying statistical power to detect correlation is well established^42^. We used equation (2) derived from Ref.^42^ to determine the smallest correlation detectable with at least 1 – *β* = 80% of statistical power at a significance level of *α* = 5%:

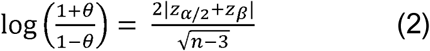

In equation (2), *n* represents the size of the sample used to estimate *θ*, and *z*_*α*/2_ and *z_β_* are the *α*/2- and *β*-upper quantiles of the standard normal distribution (mean 0 and variance 1). With *α* = 5% and *β* = 20%, *z*_*α*/2_ ≈ 1.96 and *z_β_* ≈ 0.84. We can therefore detect GPD as small as 1.2% and 0.5% in GERA and UKB respectively, and 0.4% for the meta-analysis of UKB and GERA. For the analysis of mate-pairs we can detect correlation as small as 1.5%.

### SNP Genotyping

#### UK Biobank data

We used genotyped and imputed allele counts at 1,312,100 HM3 SNPs in 487,409 participants of the UKBiobank^30,43^ (UKB). Extensive description of data can be found here^27^. We restricted the analysis to participants of European ancestry and SNPs with imputation quality ≥0.3, minor allele frequency (MAF) ≥1% and Hardy-Weinberg equilibrium test *p*-value > 10^−6^. Ancestry assignment was performed as follows. We calculated the first two principal components from 2,504 participants of the 1,000 Genomes Project^44^ with known ancestries. We then projected UKB participants onto those principal components using SNP loadings of each principal component. We assigned each individual to one of five super-populations in the 1,000 Genomes data: European, African, East Asian, South Asian and Admixed. Our algorithm calculated, for each participant, the probability of membership to the European super-population conditional on their principal components coordinates. The 456,426 out of the original 487,409 participants who had a probability of membership > 0.9 for the European cluster were assigned to the European super-population. Next, to obtain a sample of conventionally unrelated individuals, we estimated the genetic relationship matrix (GRM) for individuals in the subsample of Europeans using GCTA^31^ version 1.9. We iteratively dropped one member from each pair of individuals whose estimated relationship coefficient exceeded 0.05, until no pairs of individuals with a relationship coefficient above 0.05 remained in the sample. This restriction resulted in a sample of 348,502 conventionally unrelated Europeans. In total, we included 348,502 participants in this analysis and 1,124,803 SNPs. The North West Multi-centre Research Ethics Committee (MREC) approved the study and all participants in the UKB study analyzed here provided written informed consent.

#### Genetic Epidemiology Research on Adult Health and Aging (GERA) cohort data

We analyzed 60,586 participants of the GERA cohort using genotype data only. Ancestry was assigned using a procedure similar to that described for UKB. Genotype quality control involved standard filters (exclusion of SNPs with missing rate ≥ 0.02, Hardy-Weinberg equilibrium test *p*-value > 10^−6^ or minor allele count < 1, and removing individuals with missing rate ≥ 0.02). We imputed genotypes data to the 1,000 Genomes reference panel using IMPUTE2 software. We used GCTA to estimate the GRM of all participants using HM3 SNPs (MAF ≥ 0.01 and imputation INFO score ≥ 0.3). We finally include in the analysis 53,991 unrelated (GRM < 0.05) European participants with genotypes at 1,163,290 HM3 SNPs.

#### Health and Retirement Study (HRS)

We analysed 8,552 unrelated (GRM < 0.05) participants of the HRS cohort. GRM was calculated from 1,118,526 SNPs HM3 imputed to the 1,000 Genomes reference panel using IMPUTE2 software. SNP and samples quality control was similar to what described above for GERA.

#### SNP selection

We used the LD clumping algorithm implemented in PLINK to identify for each trait near independent SNPs (LD threshold r^2^<0.1 for SNPs < 1 Mb apart and association *p*-value < 0.005). LD clumping was performed using genotypes from HRS participants. We restricted the analysis to 1,060,523 HM3 SNPs that passed all quality controls in UKB, GERA and HRS datasets.

### Sample overlap

The Okbay *et al*. (2016) GWAS of educational attainment, the Sniekers *et al*. (2017) GWAS of intelligence quotient al. (2017) and the Kemp *et al*. (2017) GWAS of bone mineral density, included ∼150,000 participants of the UKB (first release of genotypes). To avoid bias due to sample overlap, analyses performed in UKB for these traits were restricted to 238,193 unrelated participants (UKB release 2 only), who also were not related to any of the participants included in those studies (UKB release 1). Participants of the HRS cohort were included in the Okbay *et al*. (2016) study as well as in the Day *et al*. (2015) GWAS of the onset of menopause. For the other GWAS considered in this study, no sample overlap with UKB, GERA or HRS was reported.

### Sib pairs

#### Selection

We used 21,783 sib pairs of European ancestry previously identified in the UKB^30^ using identity-by-descent sharing estimated from SNP data. We applied the within-family QFAM procedure of PLINK, as in Robinson *et al*. (2015)^27^, to assess the association between HM3 SNPs and height and EA. When applied to sib pairs, this procedure is equivalent to regressing the difference of height or EA between sibs onto the difference of allele counts. These estimates are therefore robust to population stratification. For height, we moreover performed a sample size weighted meta-analysis of estimates from the Robinson *et al*. (2015) study in 17,500 quasi-independent sib pairs, with those obtained in the UKB and used these newly estimated effect sizes to re-estimate GPD in UKB (not including any of the sib-pairs), GERA and HRS. In total we used 21,783 sib-pairs for EA and 39,283 sib-pairs for height.

#### Effective sample size

We defined the effective sample size (*N_eff_*) of within-family GWAS using *N_pairs_* independent sib pairs as the sample size of a standard GWAS (where SNP effect are estimated from simple linear regression) such that the estimated SNP effects from the within-family GWAS have the same expected sampling variance as the estimated SNP effects from standard GWAS. This leads to the following equation (derived in **Supplementary Note 4**)

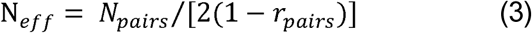

In equation (3), *r_pairs_* represents the phenotypic correlation between sibs. We observed between sibs identified in UKB a correlation ∼0.5 for height and ∼0.3 for educational attainment. Therefore, the corresponding effective sample sizes for the within-family GWAS of height and EA are 39,283/(2×(1-0.5)) = 39,283 and 21,283/(2×(1-0.3)) = 15,559.

### Mate pairs

We first used 999 unique mate pairs from 1,065 trios composed of both parents and one child, identified among UKB participants using identity-by-descent sharing estimated from SNP data. Details about software and algorithms used to identify those trios are given in Ref.^30^ To increase power, we also used household sharing information to identify putative spouse pairs among UKB participants with European ancestry. We therefore selected 18,984 (including 117 from the trios) sex-discordant pairs of participants, recruited from the same centre, who reported living with their spouse or partner in the same type of accommodation, at the same location (east and north coordinates rounded to 1 kilometre), for the same amount of time, with the same number of people in the household, with the same household income, with the same number of smoker in the household, with the same Townsend deprivation index and with a genetic relationship < 0.05.

### Data availability

We used genotypic data from the Resource for Genetic Epidemiology Research on Adult Health and Aging (GERA: dbGaP phs000674.v2.p2), genotypic and phenotypic from the Health and Retirement Study (HRS: dbGaP phs000428.v1.p1), as well as genotypic and phenotypic data from the UK Biobank under project 12505.

## Acknowledgements

This research was supported by the Australian Research Council (DP130102666, DP160103860, DP160102400), the Australian National Health and Medical Research Council (1078037, 1078901, 1103418, 1107258, 1127440 and 1113400), the National Institute of Health (NIH grants R01AG042568, P01GM099568 and R01MH100141), the Sylvia & Charles Viertel Charitable Foundation. The Genetic Epidemiology Research on Adult Health and Aging study was supported by grant RC2 AG036607 from the National Institutes of Health, grants from the Robert Wood Johnson Foundation, the Ellison Medical Foundation, the Wayne and Gladys Valley Foundation and Kaiser Permanente. The authors thank the Kaiser Permanente Medical Care Plan, Northern California Region (KPNC) members who have generously agreed to participate in the Kaiser Permanente Research Program on Genes, Environment and Health (RPGEH). This research has been conducted using the UK Biobank Resource under project 12505. We thank Bill Hill for helpful comments and suggestions on the manuscript.

## Author contributions

P.M.V, L.Y., M.R.R, J.Y. and M.E.G. conceived and designed the study. L.Y., M.T. and N.W. curated summary statistics. L.Y. and P.M.V derived the theory. Y.Y., M.T., J.G., K.E.K and L.Y. performed mate pairs analyses. M.C.K., P.T., D.B. and D.C. helped developing the methodology and interpret the results. P.M.V., N.W., M.R.R. and L.Y. performed sib-pairs analyses. K.E.K. and L.Y. performed quality control of UKB data. L.Y. and M.R.R. performed statistical analyses and simulations. L.Y. and P.M.V wrote the manuscript with the participation of all authors.

**Fig. S1.**
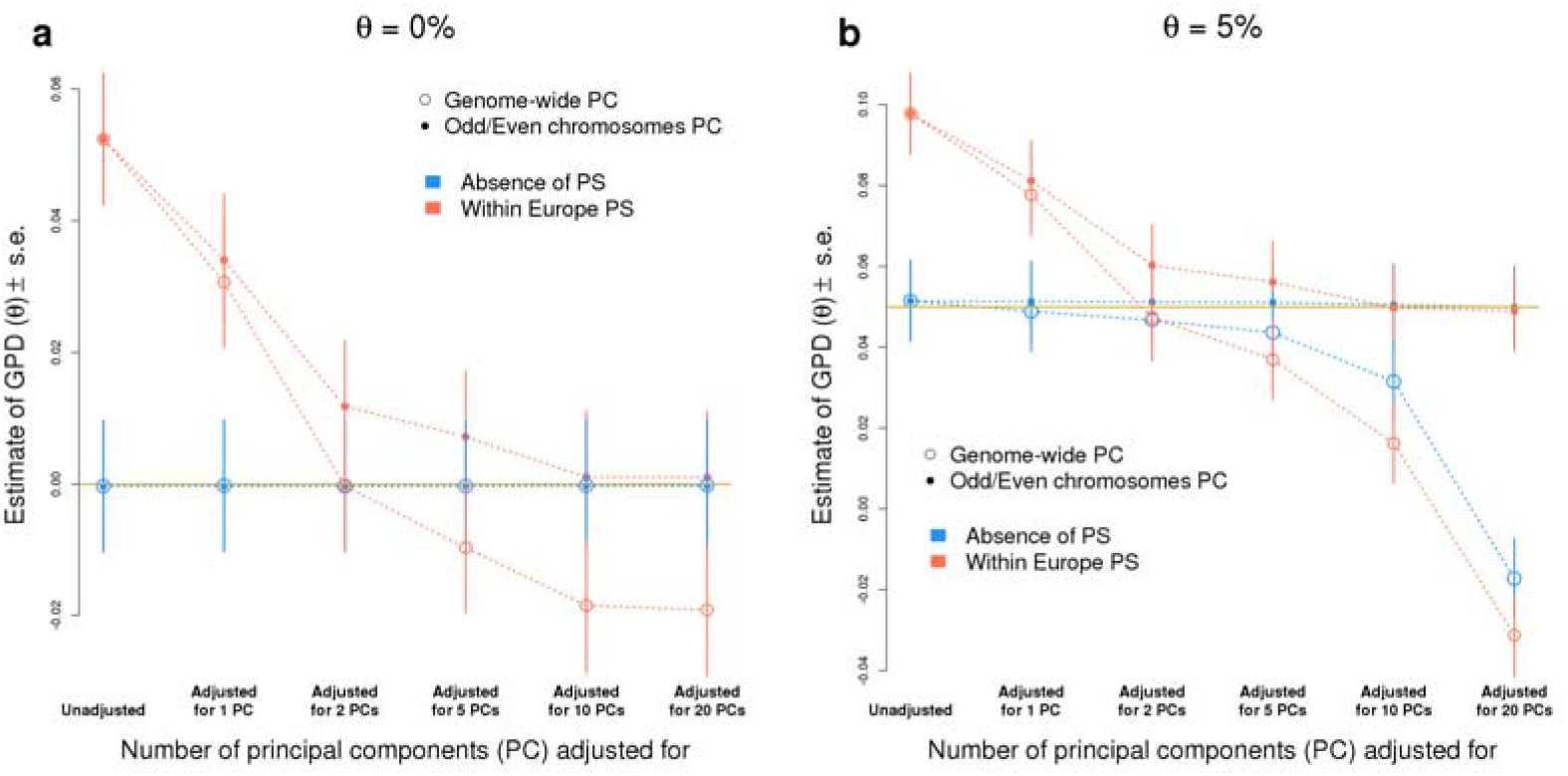
Estimates of gametic phase disequilibrium (GPD), in simulated data (N=10,000) using allele frequencies of 697 height-associated SNPs, as a function of the number of genotypic principal components adjusted to correct for population stratification. Data were simulated assuming either no population stratification or within-Europe population stratification (**Supplementary Note 2**). In both cases, data were simulated under pure random mating (*θ*=0, panel **a**) or under assortative mating (*θ*=5%, panel **b**). Estimates are obtained from unadjusted linear regression or adjusted for 1, 2, 5, 10 and 20 first principal components. Principal components were either calculated from SNPs on odd and even chromosomes (genome-wide principal components) or from SNPs on odd or even chromosomes separately. Standard Error (s.e.).

**Fig. S2.**
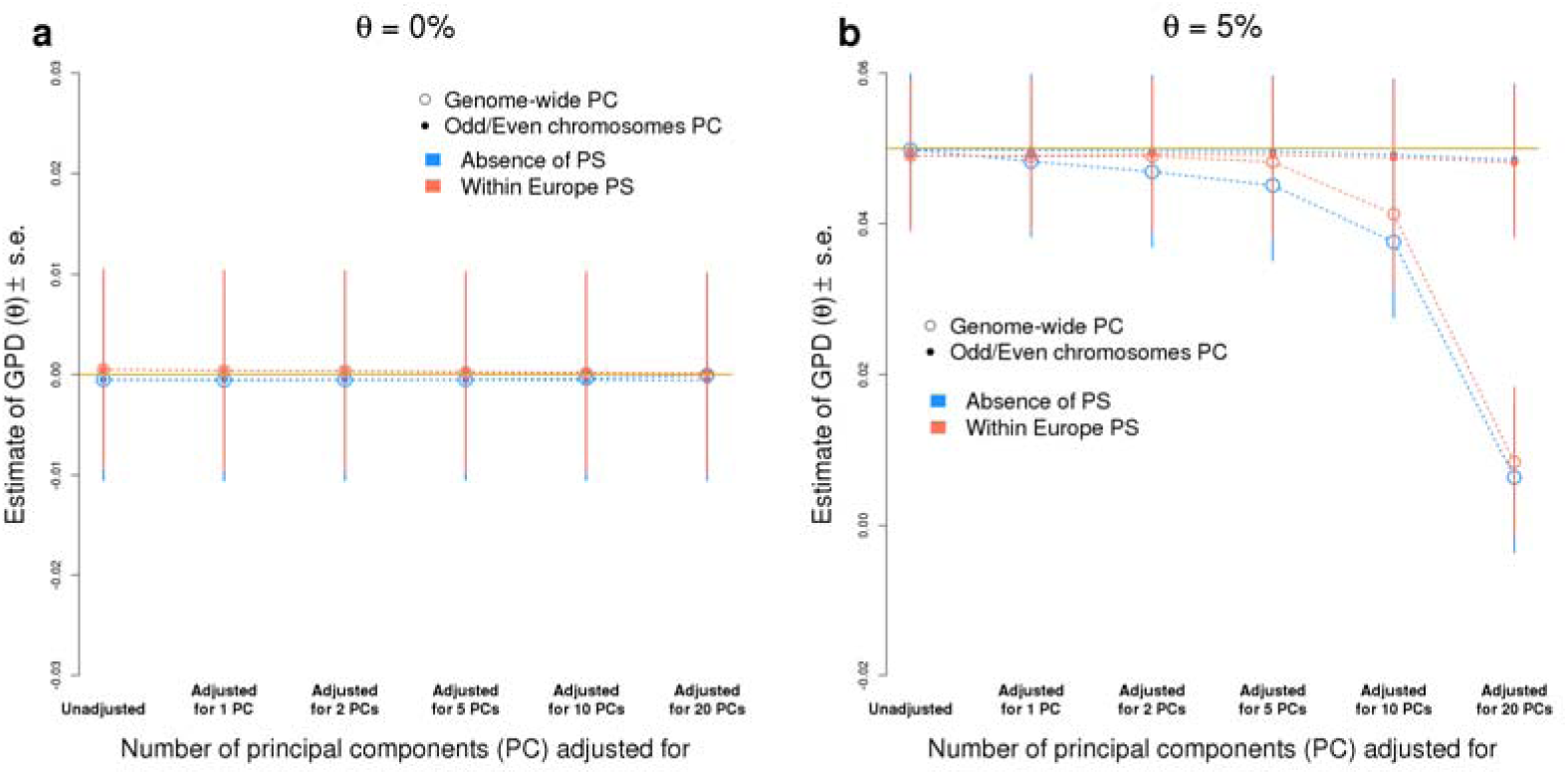
Estimates of gametic phase disequilibrium (GPD), in simulated data (N=10,000) using allele frequencies of 1,000 randomly selected Hap Map 3 SNPs, as a function of the number of genotypic principal components adjusted to correct for population stratification. Data were simulated assuming either no population stratification or within-Europe population stratification (**Supplementary Note 2**). In both cases, data were simulated under pure random mating (*θ*=0, panel **a**) or under assortative mating (*θ*=5%, panel **b**). Estimates are obtained from unadjusted linear regression or adjusted for 1, 2, 5, 10 and 20 first principal components. Principal components were either calculated from SNPs on odd and even chromosomes (genome-wide principal components) or from SNPs on odd or even chromosomes separately. Standard Error (s.e.).

**Fig. S3.**
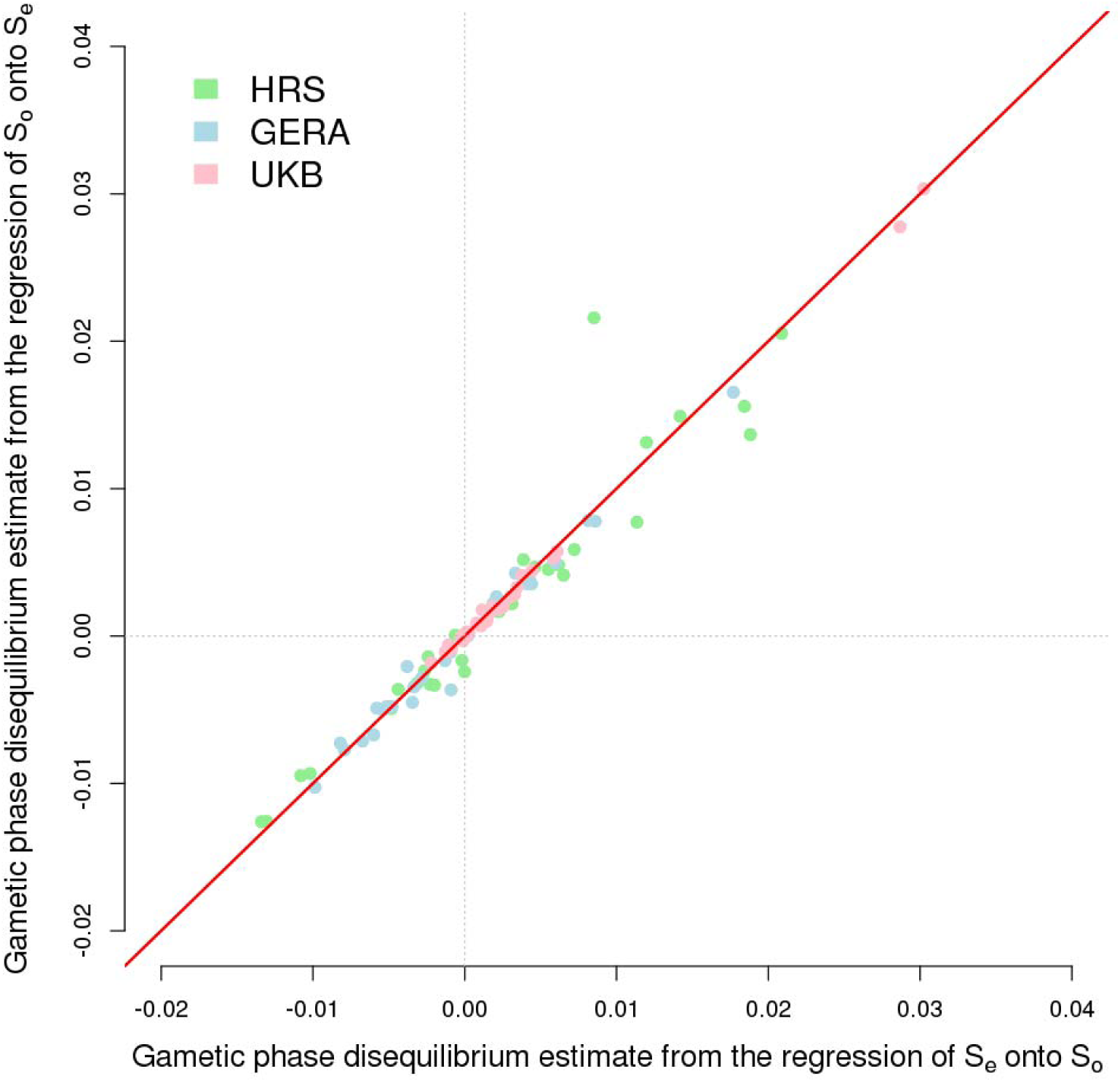
Comparison of estimates of gametic phase disequilibrium (GPD) from two approaches: (a) from the regression of genetic predictors from SNPs on even chromosomes (S_e_) onto of genetic predictors from SNPs on odd chromosomes (S_o_); and (b) from the regression of S_o_ onto S_e_. The x-axis corresponds to approach (a) and y-axis to approach (b). The correlation between these two estimators is ∼0.99 across all three cohorts UKB (N=348,502), GERA (N=53,991) and HRS (N=8,552).

**Table S1.** Within-individual correlation (*θ* in %) between trait-specific genetic predictors from SNPs on odd chromosomes versus even chromosomes for 32 traits and diseases. Estimates from UKB (N=348,502), GERA (N=53,991) and HRS (N=8,552) are denoted *θ*_UKB_, *θ_GERA_* and *θ*_HRS_ respectively; and estimate from the meta-analysis of these three estimates is denoted *θ*_META_. ^∗^As the HRS cohort was included in the educational attainment GWAS and the age at menopause GWAS, *θ*_META_ calculations for these traits did not include data from HRS. All estimates are adjusted for 20 genotypic principal components from SNPs on either odd or even chromosomes. Standard errors (s.e.) of estimated correlations within each cohort are approximately inversely proportional to the square-root of the sample size (N). For UKB s.e. ∼0.17%, for GERA s.e.∼0.43% and for HRS s.e.∼1.1%.

| Traits/Diseases | Number of SNPs<br>to estimate $\theta$ | $\theta_{\text{UKB}}$ | $\theta_{\text{GERA}}$ | $\theta_{\text{HRS}}$ | $\theta_{\text{META}}$ | Standard Error<br>($\theta_{\text{META}}$) | Meta-analysis<br>$p$-value | Heterogeneity<br>$p$-value |
| --- | --- | --- | --- | --- | --- | --- | --- | --- |
| Height | 9,447 | 3.02 | 4.15 | 4.44 | 3.20 | 0.16 | $6.5 \times 10^{-89}$ | 0.03 |
| Number of years of education | 4,618 | 2.87 | 1.77 | 8.95* | 2.67 | 0.19 | $6 \times 10^{-46}$ | 0.02 |
| Intelligence Quotient | 3,356 | 0.61 | 0.33 | 1.42 | 0.58 | 0.19 | $2.3 \times 10^{-3}$ | 0.65 |
| Former smoker | 2,011 | 0.44 | 0.82 | -1.02 | 0.46 | 0.16 | $3.8 \times 10^{-3}$ | 0.30 |
| Attention Deficit Hyperactivity<br>Disorder | 2,902 | 0.58 | -0.79 | -0.31 | 0.38 | 0.16 | 0.02 | 0.01 |
| Age at menopause | 2,263 | 0.33 | 0.41 | 4.49* | 0.34 | 0.16 | 0.03 | 0.85 |
| Alcohol consumption<br>(Continuous measurement) | 2,186 | 0.19 | 0.86 | 0.31 | 0.28 | 0.17 | 0.09 | 0.39 |
| Bipolar Disorder | 2,358 | 0.38 | -0.58 | 0.39 | 0.25 | 0.16 | 0.13 | 0.14 |
| Age at menarche | 3,385 | 0.18 | 0.38 | 1.56 | 0.24 | 0.16 | 0.13 | 0.42 |
| Systolic Blood Pressure | 1,687 | 0.22 | 0.44 | -0.24 | 0.24 | 0.17 | 0.17 | 0.84 |
| Crohn's Disease | 2,669 | 0.14 | 0.60 | 1.20 | 0.22 | 0.16 | 0.17 | 0.45 |
| Extraversion | 1,666 | 0.30 | -0.51 | -1.30 | 0.16 | 0.16 | 0.33 | 0.10 |
| Ever smoked | 2,281 | 0.25 | -0.60 | 0.72 | 0.15 | 0.16 | 0.35 | 0.18 |
| Major Depressive Disorder | 1,958 | -0.11 | -0.82 | 1.88 | -0.16 | 0.18 | 0.35 | 0.09 |
| Rheumatoid Arthritis | 2,162 | 0.11 | -0.09 | 0.85 | 0.10 | 0.11 | 0.38 | 0.47 |
| Openness | 1,674 | -0.11 | -0.54 | 2.09 | -0.12 | 0.16 | 0.43 | 0.08 |
| Type 2 Diabetes | 2,904 | 0.18 | -0.17 | -0.33 | 0.12 | 0.16 | 0.46 | 0.72 |
| Agreeableness | 1,651 | -0.01 | -0.67 | -0.44 | -0.11 | 0.16 | 0.49 | 0.34 |
| Anxiety case / control | 1,872 | -0.18 | 0.37 | 0.45 | -0.10 | 0.14 | 0.49 | 0.38 |
| Number of Cigarettes per Day | 1,947 | -0.13 | 0.02 | -0.26 | -0.11 | 0.17 | 0.52 | 0.95 |
| Conscientiousness | 1,688 | 0.16 | -0.28 | -0.06 | 0.10 | 0.16 | 0.55 | 0.63 |
| Coronary Artery Disease | 2,158 | 0.09 | 0.22 | -1.26 | 0.08 | 0.16 | 0.63 | 0.44 |
| Bone Mineral Density | 5,859 | 0.34 | -0.99 | 0.00 | 0.09 | 0.19 | 0.64 | 0.03 |
| Autism Spectrum Disorder | 1,811 | 0.11 | -0.11 | -0.17 | 0.07 | 0.16 | 0.65 | 0.88 |
| Age at smoking onset | 1,925 | -0.03 | -0.34 | -0.23 | -0.07 | 0.17 | 0.67 | 0.82 |
| Anxiety (Factor score) | 2,005 | -0.09 | 0.01 | 0.46 | -0.06 | 0.16 | 0.68 | 0.87 |
| Post-Traumatic Stress Disorder | 1,814 | -0.09 | 0.19 | 0.22 | -0.05 | 0.16 | 0.77 | 0.82 |
| Body Mass Index (BMI) | 2,362 | 0.01 | -0.48 | 0.65 | -0.04 | 0.15 | 0.78 | 0.43 |
| Diastolic Blood Pressure | 1,663 | -0.01 | 0.58 | -1.08 | 0.05 | 0.17 | 0.78 | 0.31 |
| Waist/Hip ratio (adj. BMI) | 1,940 | 0.01 | 0.27 | -0.49 | 0.03 | 0.16 | 0.85 | 0.77 |
| Alcohol consumption<br>(Dichotomous measurement) | 1,614 | 0.00 | -0.38 | 1.13 | -0.03 | 0.19 | 0.87 | 0.56 |
| Schizophrenia | 5,617 | 0.03 | -0.34 | 0.62 | -0.01 | 0.15 | 0.94 | 0.59 |

**Table S2.** Between mates correlation (in %) of genetic predictors of 32 traits and diseases. Estimates are adjusted for 20 genotypic principal components from SNPs on both odd and even chromosomes. Significant correlations (*p*-value < 0.05/30) are marked with a “^∗^” Pairs involving UKB participants included in the EA, IQ and BMD GWAS were removed.

| **Traits/Diseases** | $r_m$ (%) | s.e. ($r_m$) | $p$-value | Number of<br>mate pairs |
| --- | --- | --- | --- | --- |
| **Height\*** | **5.87** | **0.79** | **$9.8 \times 10^{-14}$** | **18,984** |
| **Number of years of education (EA)\*** | **6.13** | **0.94** | **$7.3 \times 10^{-11}$** | **10,854** |
| Intelligence Quotient (IQ) | 1.08 | 0.96 | 0.26 | 10,854 |
| Former smoker | -0.51 | 0.74 | 0.48 | 18,984 |
| Attention Deficit Hyperactivity Disorder | 0.25 | 0.73 | 0.73 | 18,984 |
| Age at menopause | -0.23 | 0.73 | 0.75 | 18,984 |
| Alcohol consumption<br>(Continuous measurement) | -0.49 | 0.73 | 0.5 | 18,984 |
| Bipolar Disorder | 0.23 | 0.72 | 0.75 | 18,984 |
| Age at menarche | 0.32 | 0.72 | 0.66 | 18,984 |
| Systolic Blood Pressure | 0.56 | 0.72 | 0.44 | 18,984 |
| Crohn's Disease | 0.85 | 0.73 | 0.24 | 18,984 |
| Extraversion | -1.38 | 0.72 | 0.06 | 18,984 |
| Ever smoked | 0.68 | 0.74 | 0.36 | 18,984 |
| Major Depressive Disorder | 0.34 | 0.73 | 0.64 | 18,984 |
| Rheumatoid Arthritis | 1.10 | 0.75 | 0.14 | 18,984 |
| Openness | -0.12 | 0.73 | 0.87 | 18,984 |
| Type 2 Diabetes | 1.36 | 0.73 | 0.06 | 18,984 |
| Agreeableness | -0.75 | 0.72 | 0.29 | 18,984 |
| Anxiety case / control | 0.88 | 0.72 | 0.22 | 18,984 |
| Number of Cigarettes per Day | 1.20 | 0.72 | 0.96 | 18,984 |
| Conscientiousness | 0.08 | 0.72 | 0.91 | 18,984 |
| Coronary Artery Disease | 0.07 | 0.72 | 0.93 | 18,984 |
| Bone Mineral Density (BMD) | -1.02 | 1.00 | 0.3 | 10,854 |
| Autism Spectrum Disorder | 2.15 | 0.73 | 0.003 | 18,984 |
| Age at smoking onset | -0.36 | 0.73 | 0.62 | 18,984 |
| Anxiety (Factor score) | 0.27 | 0.72 | 0.71 | 18,984 |
| Post-Traumatic Stress Disorder | 0.58 | 0.72 | 0.42 | 18,984 |
| Body Mass Index (BMI) | 0.73 | 0.72 | 0.3 | 18,984 |
| Diastolic Blood Pressure | -0.56 | 0.73 | 0.44 | 18,984 |
| Waist/Hip ratio (adj. BMI) | 0.10 | 0.73 | 0.89 | 18,984 |
| Alcohol consumption<br>(Dichotomous measurement) | 1.21 | 0.72 | 0.09 | 18,984 |
| Schizophrenia | 2.05 | 0.76 | 0.006 | 18,984 |

**Table S3.** Description of summary statistics from genome-wide associations (GWAS) on 32 traits analysed in this study. LD score regression (LDSC) was applied to each set of summary statistics to calculate SNP heritability and LDSC ratio statistics, a measure of population stratification. The LDSC software (version 1.0) does not calculate standard errors (s.e.) of the ratio statistic when it is < 0. In these cases, s.e. of the ratio statistics were replaced with “.”. URLs for downloading summary statistics used in this study are given in **Supplementary Note 1**.

| **Traits/Diseases** | **GWAS**<br>**median**<br>**sample size** | **LDSC**<br>**heritability**<br>**(h<sup>2</sup>)** | **LDSC**<br>**heritability**<br>**(standard**<br>**error)** | **LDSC**<br>**intercept** | **LDSC**<br>**intercept**<br>**(standard**<br>**error)** | **LDSC Ratio**<br>**statistic** | **LDSC**<br>**ratio**<br>**statistic**<br>**(standard**<br>**error)** |
| --- | --- | --- | --- | --- | --- | --- | --- |
| Number of years of education | 293,723 | 12.7% | 0.4% | 0.93 | 0.01 | -0.11 | . |
| Intelligence Quotient | 78,308 | 18.8% | 1.0% | 1.01 | 0.01 | 0.03 | 0.03 |
| Height | 252,083 | 34.2% | 1.8% | 1.23 | 0.03 | 0.12 | 0.02 |
| Body Mass Index | 233,692 | 13.5% | 0.7% | 0.66 | 0.01 | -1.30 | . |
| Waist/Hip ratio (adj. BMI) | 142,438 | 9.0% | 0.7% | 0.86 | 0.01 | -1.44 | . |
| Attention Deficit Hyperactivity<br>Disorder | 55,374 | 23.2% | 1.5% | 1.03 | 0.01 | 0.10 | 0.03 |
| Autism Spectrum Disorder | 13,574 | 33.5% | 4.4% | 0.99 | 0.01 | -0.17 | . |
| Bipolar Disorder | 16,731 | 47.1% | 4.1% | 1.01 | 0.01 | 0.03 | 0.06 |
| Anxiety case / control | 17,310 | 7.6% | 3.0% | 1.00 | 0.01 | 0.10 | 0.25 |
| Anxiety (Factor score) | 18,186 | 7.0% | 2.7% | 1.00 | 0.01 | -0.05 | . |
| Major Depressive Disorder | 18,759 | 17.4% | 2.6% | 1.00 | 0.01 | 0.05 | 0.09 |
| Post-Traumatic Stress Disorder | 53,293 | 1.8% | 0.8% | 0.99 | 0.01 | -0.48 | . |
| Schizophrenia (SCZ) | 150,064 | 24.3% | 1.0% | 1.04 | 0.01 | 0.05 | 0.02 |
| Coronary Artery Disease | 184,305 | 6.7% | 0.5% | 0.88 | 0.01 | -0.88 | . |
| Type 2 Diabetes (T2D) | 152,599 | 7.8% | 0.6% | 1.00 | 0.01 | -0.01 | . |
| Crohn's Disease | 20,883 | 50.7% | 5.8% | 1.02 | 0.01 | 0.09 | 0.05 |
| Rheumatoid Arthritis | 58,284 | 15.0% | 2.6% | 0.94 | 0.01 | -0.50 | . |
| Systolic Blood Pressure (SBP) | 67,211 | 11.2% | 0.9% | 0.90 | 0.01 | -1.77 | . |
| Diastolic Blood Pressure (DBP) | 67,201 | 11.0% | 1.0% | 0.91 | 0.01 | -1.61 | . |
| Age at menopause | 70,000 | 13.4% | 1.6% | 0.99 | 0.01 | -0.06 | . |
| Age at menarche | 182,416 | 11.9% | 0.6% | 0.97 | 0.01 | -0.07 | . |
| Conscientiousness | 17,375 | 7.5% | 3.2% | 1.00 | 0.01 | -0.06 | . |
| Agreeableness | 17,375 | 1.7% | 3.0% | 1.00 | 0.01 | 0.32 | 0.97 |
| Openness | 17,375 | 12.2% | 2.8% | 0.99 | 0.01 | -0.40 | . |
| Extraversion | 17,375 | 4.5% | 2.7% | 1.00 | 0.01 | -0.11 | . |
| Ever smoked | 74,053 | 7.5% | 0.7% | 1.00 | 0.01 | -0.03 | . |
| Age at smoking onset | 74,053 | 2.0% | 0.6% | 1.00 | 0.01 | -0.04 | . |
| Number of Cigarettes per Day | 74,053 | 3.1% | 0.8% | 1.00 | 0.01 | 0.09 | 0.12 |
| Former smoker | 74,053 | 3.5% | 0.6% | 1.00 | 0.01 | 0.01 | 0.13 |
| Alcohol consumption (Binary) | 70,460 | 0.8% | 0.6% | 0.99 | 0.01 | -3.52 | . |
| Alcohol consumption Continuous | 70,460 | 5.2% | 0.8% | 1.01 | 0.01 | 0.17 | 0.09 |
| Bone Mineral Density (BMD) | 142,487 | 37.6% | 4.1% | 1.08 | 0.03 | 0.07 | 0.03 |

